# Clogging-free continuous operation with whole blood in a radial pillar device (RAPID)

**DOI:** 10.1101/197749

**Authors:** Ninad Mehendale, Oshin Sharma, Shilpi Pandey, Debjani Paul

## Abstract

Pillar-based passive microfluidic devices combine the advantages of simple designs, low device footprint, and high selectivity for size-based separation of blood cells. Most of these device designs have been validated with dilute blood samples. Handling whole blood in pillar-based devices is extremely challenging due to clogging. The high proportion of cells (particularly red blood cells) in blood, the varying sizes and stiffness of the different blood cells, and the tendency of the cells to aggregate lead to clogging of the pillars within a short period. We recently reported a <u>ra</u>dial <u>pi</u>llar <u>d</u>evice (RAPID) design for continuous and high throughput separation of multi-sized rigid polystyrene particles in a single experiment. In this manuscript, we have given detailed guidelines to modify the design of RAPID for any application with deformable objects (e.g. cells). We have adapted RAPID to work with blood samples directly without any pre-processing steps. We were successful in operating the device with whole blood for almost 6 hours, which is difficult to achieve with most pillar-based devices. Finally, we demonstrated up to ~ 60-fold enrichment of platelets as an illustration of the improved device design. Whole blood pillar-based platelet clog-free RAPID

## 1 Introduction

Blood analysis can provide critical insights into the general health and immune system of an individual. Therefore, one of the most common tests prescribed at the first point of diagnosis is the blood test. Tests on whole blood samples are also performed during clinical research, drug discovery, and the development of new diagnostic technologies. Many of the tests involving plasma, platelets, white blood cells (WBCs) or rare cells (e.g. circulating tumor cells, fetal cells, etc.) are affected by the presence of the red blood cells (RBCs). This is because RBCs outnumber platelets (20:1) and WBCs (500:1) [1]. A sample preparation step is often required to remove or lyse RBCs prior to analysis. As discussed by Mariella [2], integrating an on-chip sample preparation step still remains the ‘weak link’ in the development of microfluidic devices. Hence, there is a need for microfluidic devices that can handle whole blood and separate its different components. One of the simplest and highly reported label-free microfluidic techniques for separating cells is based on size exclusion [3, 4, 5, 6, 7, 8, 9]. This is achieved by fabricating a network of pillars (or pores), where the pillar gaps (or the pore sizes) are comparable to the size of a single cell. This technique has been adapted to sort WBCs (nucleated stiff cells with 10 - 30 *μ*m diameter), RBCs (non-nucleated and deformable biconcave cells with 6 - 8 *μ*m diameter and 2 - 3 *μ*m thickness) and platelets (stiff cell fragments that are 2 - 3 *μ*m in size). The operation of most of these devices was demonstrated with dilute blood [10]. Diluting the blood has two disadvantages: (1) it leads to an additional off-chip processing step; and (2) it reduces the concentration of the target analyte for downstream applications. Dilute samples are used because pillar-based designs suffer from a fundamental disadvantage of clogging when operated with concentrated cell suspensions or whole blood. As a result, the device becomes unusable. Some remedial measures to free the clogged cells include making the pillar gaps ratchet-shaped [11], oscillating the fluid flow [12], perfusion [9], etc. Cross-flow filters were explicitly designed to avoid clogging [13, 10, 14, 15, 16]. In this design, many of the smaller cells remain in the main flow, thereby affecting the selectivity of separation. The large footprint of cross-flow devices also leads to sample volume loss. Deterministic lateral displacement devices can perform size-based cell separation with very high resolutions, but cannot handle concentrated samples due to enhanced particle-post and particle-particle interactions [17, 18, 19].

There are some reports on the use of whole blood, primarily for plasma separation [20, 21], in pillar-based microfluidic devices. These devices have reported operation times of the order of a few minutes using a few *μ*l of blood [7, 6, 21, 8]. Some of them have also reported clogging and hemolysis [22]. It is not clear whether these devices are capable of longer continuous operation and handling larger sample volumes. As discussed by Kersaudy-Kerhoas and Sollier [22], continuous operation is needed for diverse applications such as studying the effect of drugs in real-time, cell migration, etc. Moreover, the capture of rare cells or biomolecules from blood requires the microfluidic device to handle large (~ ml) sample volumes [23, 24, 25].

Recently we developed a passive <u>ra</u>dial <u>pi</u>llar <u>d</u>evice (RAPID) that can operate in a continuous and clog-free manner [26]. We demonstrated its operation by separating a mixture of polystyrene particles with high purity, throughput, and recovery. There are specific design challenges when one works with deformable samples, such as RBCs. Therefore, in this report, we adopted the design of RAPID to work continuously with whole blood and operated it for close to 6 h. As an illustration of the improved design, we also achieved up to ~ 60X platelet enrichment without any sample pre-processing.

## 2 Materials and Methods

### 2.1 Equipment and chemicals

SU-8 2005 and its developer were obtained from MicroChem Corporation (Westborough, USA). Sylgard 184 (PDMS) was purchased from Dow Corning Corporation (Michigan, USA). Common chemicals, such as, ethanol, isopropyl alcohol, sodium hypochlorite, etc. were obtained from Thomas Baker (Mumbai, India) and used without further purification. Normal saline (0.9% NaCl) was bought locally. 60mm × 24 mm glass cover slips (No. 1) were bought from Blue Star, Mumbai, India. Microfluidic connectors (barb-to-barb WW-30626-48 and luer-to-barb WW-30800-06) from Cole Parmer (Mumbai, India) were used, while 1.5 mm diameter Tygon tubing (formulation 2375) was used for chip-to-syringe connections. We used 1 ml plastic syringes from Becton-Dickinson (Mumbai, India). BD vacutainer tubes coated with K2-EDTA were bought from Fisher scientific, USA, for storing blood samples.

Spin coating of photoresist was carried out on model WS-400BZ from Laurell Technologies Corporation (PA, USA), and UV exposure was done in a MJB4 mask aligner from Karl Suss. The height of the pillars was measured using an Ambios XP2 profilometer. PDMS-glass bonding was performed using a Harrick plasma cleaner (PDC 32G). A syringe pump (model 111, Cole Parmer) was used to control the blood flow. A vibrator motor (PNN7RB55PW2) was bought locally and attached to the tubes carrying blood. Platelet counts were performed using a hematology analyzer (Sysmax XS-800i). Images of the device were acquired using a Nikon Eclipse Ti inverted microscope fitted with a 40X (1.3 NA) objective.

### 2.2 Microfluidic chip fabrication

RAPID was fabricated in PDMS (Sylgard 184) using standard soft lithography. A 2-inch diameter silicon wafer was cleaned by RCA technique and dehydrated on a hot plate at 120 *°*C for 20 min. SU-8 2005 photoresist was spin-coated on the silicon wafer (500 rpm for 15 sec, followed by 3000 rpm for 30 sec). The resist was pre-baked on a hot plate at 90*°*C for 3 min, followed by UV exposure for 8 sec at 100 mJ/*cm*^2^. Post-exposure bake was carried out at 90*°*C for 1 min, and the pattern was developed in the SU-8 developer for 4 min. The developed pattern was rinsed in IPA and blow-dried using a nitrogen gun. Finally, a hard bake step was performed at 120*°*C for 10 min to generate the molding template. The height of the pattern was 5 *μ*m, as confirmed by profilometry (supplementary figure S1).

PDMS base and curing agent were mixed in the ratio of 10:1, degassed and poured on the SU-8 master. PDMS was cured in a hot air oven at 65*°*C for 45 min. The PDMS chip was then cut using a surgical blade and gently peeled from the SU-8 master. The RBC outlet was punched using a 26G syringe needle under a 4X microscope objective. The inlet and the platelet outlet were punched using a 1.2 mm biopsy punch. Next, the chip was bonded to a glass coverslip using plasma bonding. We used oxygen plasma for 90 sec. To strengthen the bond, the bonded chip was again placed in the oven for 30 min at 100 *°*C.

As showed in the supplementary figure S2, we reinforced the connectors and the tubing prior to handling whole blood. Two 18G (1.2 mm diameter) blunt syringe needles were connected to the inlet and the platelet outlet respectively of the bonded chip. A 26G (0.45 mm diameter) blunt needle was connected to the RBC outlet. In order to resolve the problem of device failure, the bonded chip was placed inside a 25 mm X 25 mm X 25 mm 3-D printed part. PDMS was then poured on the chip and the connectors to a height of 20 mm. The entire 3D-printed part (with the chip and the connectors) was then degassed and cured in the oven for 30 min at 100 °C. The curing temperature was chosen to be high to allow PDMS to cure quickly and prevent from getting inside the chip. Finally, the entire PDMS mold was taken out of the 3-D printed part. Luer-to-barb connectors were attached to the needles, and Tygon tubing was connected prior to commencing the experiment.

### 2.3 Preparation of blood samples

A trained phlebotomist drew 2 ml of blood from healthy volunteers after obtaining informed consent and transferred the blood to vacutainer tubes coated with K2-EDTA. The blood was used no later than 24 hours for platelet enrichment experiments.

### 2.4 Platelet enrichment experiments

A 1 ml syringe was filled with blood and mounted on a syringe pump as shown in Fig 1C. The blood inlet in the chip was connected to the syringe using the connection strategy described in the previous section. A vibrator motor was attached to the inlet tubing, midway between the chip and the syringe. Since a significant number of RBCs settle in the tube in 20 min, we developed a controller circuit to turn on the motor for 40 sec after every 20 min. The details of the circuit are given in the supplementary information.

Inlet flow rates of 600 nl/min, 700 nl/min, 1 *μ*l/min, 3 *μ*l/min, 5 *μ*l/min and 10 *μ*l/min were explored to study the effect of flow rate on different performance measures. The samples from the platelet outlet and the RBC outlet were collected in two Eppendorf tubes. The experiment was stopped when ~ 50 *μ*l of the sample was collected from both outlets. The number of WBCs, RBCs, and platelets present at the inlet and collected from the outlets (RBC and platelet) of RAPID were counted using a hematology analyzer for each flow rate. Experiments with eighteen devices were performed (N = 3 for whole blood at each of the six flow rates) to account for any device-to-device variability.

### 2.5 Image analysis

Time-lapse images (for 6h and 4h) of experiments with whole blood were recorded using an inverted microscope (Nikon Eclipse Ti) at flow rates of 2 *μ*l/min and 600 nl/min respectively. The images were acquired every 4 sec. The cell tracks were obtained from the videos using a MATLAB code. Frames were extracted and converted to grayscale images. We applied ‘speeded up robust features’(SURF) algorithm of MATLAB to the images to locate the cells. Images were corrected for orientation and scale. The difference between consecutive frames was computed to find the displacement of cells between the frames. The difference frame was converted to a binary image with a specific threshold of 10%. All noise was removed using an area filter (20 pixels). The major axis and orientation of each object in every difference image were computed. All cells within ±10*°* of the vertical were considered to be flowing radially, while objects within ±10*°* of the horizontal were considered to be in the cross flow. The velocity was calculated every 15 min.

**Figure 1:**
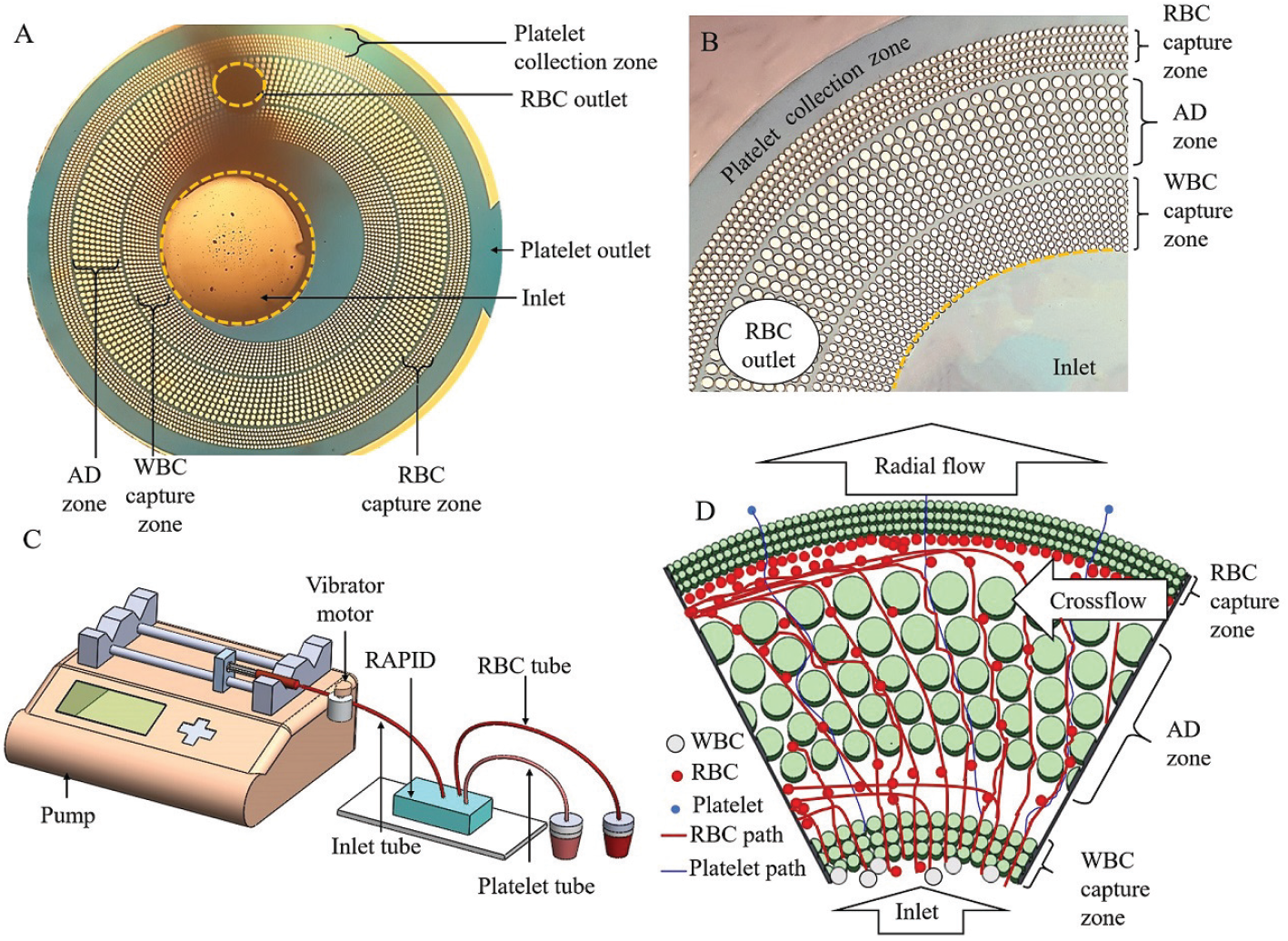
Working principle of RAPID. (A) A microscope image of RAPID, acquired at 4X magnification, shows a central inlet for introducing blood, an RBC outlet, and a platelet outlet. The pillars are arranged in concentric circles in three zones. The pillars near the inlet block WBCs and large cell aggregates, while allowing RBCs and platelets to go through. The AD zone is used to remove the RBCs along a cross-flow towards the RBC outlet. The platelets move radially outwards to the platelet collection zone. (B) A magnified image of a sector of the device shows the varying sizes of the pillars and the pillar gaps in each of the three zones. The pillar gaps are progressively decreased to separate cells according to their size and deformability. (C) The schematic diagram of the experimental set-up for platelet enrichment from whole blood is shown. It consists of a syringe pump, a vibrator motor attached to the inlet tubing, the RAPID chip and two Eppendorf tubes for collecting the RBCs and the enriched platelet solution. (D) The schematic diagram shows the radial, and the cross flows in RAPID. Most platelets follow the radial path (blue lines). The RBCs (red lines) take the radial path through the WBC capture zone, and then follow the cross flow.

## 3 Results and discussions

### 3.1 How to design RAPID?

RAPID (Fig. 1A) comprises of pillars arranged in concentric circles over three zones around a central inlet. The sample is introduced into the chip through the inlet and pumped in a radial direction. The innermost zone (zone 1) captures WBCs. The middle zone (zone 2) sets up a cross-flow (Fig. 1D) in the device. The outermost zone (zone 3) stops the RBCs and lets the platelets through.

**Figure 2:**
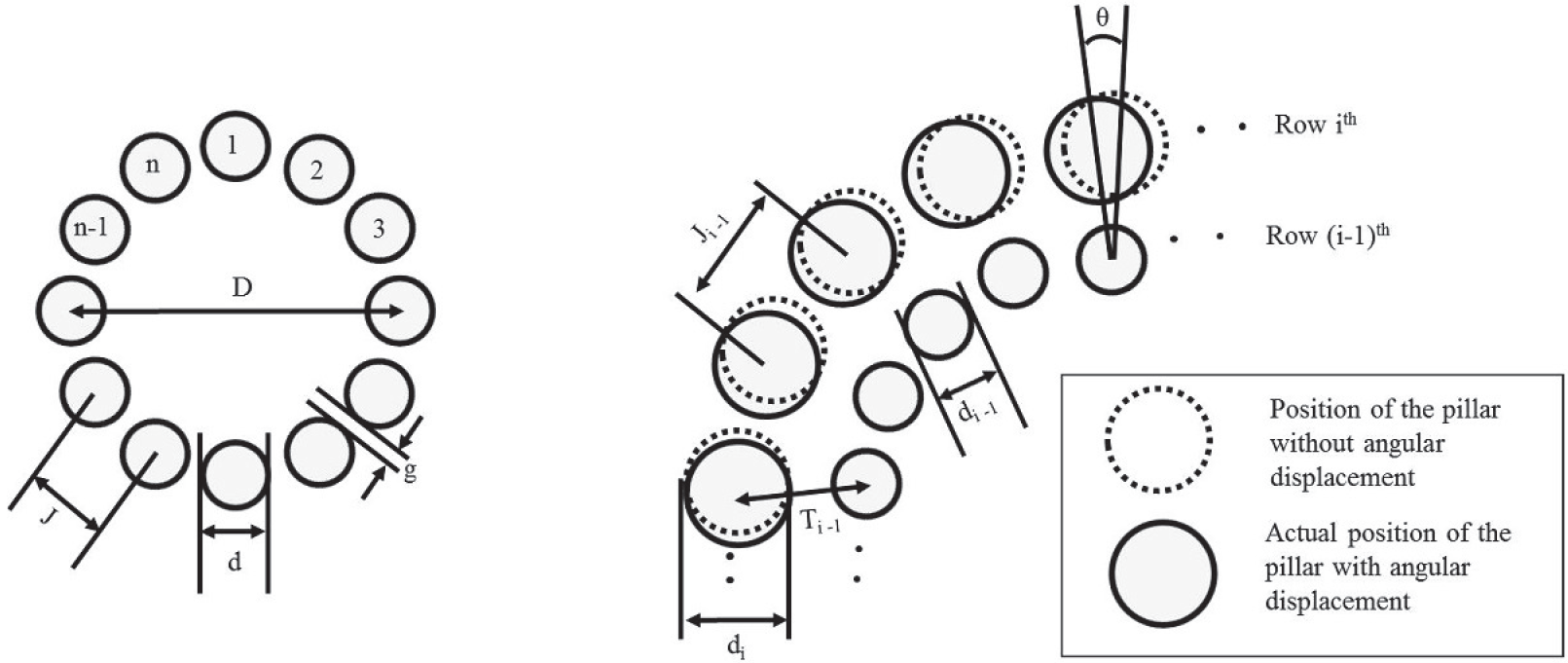
The relationship between different geometrical parameters of RAPID. (A) The scheme for choosing the pillar diameter (*d*), the pillar gap (*g*), and the number of pillars (*n*) of any particular row of RAPID. *D* is twice the distance from the center of the device to the center of the pillar. (B) The scheme to determine the angular displacement *θ* between the pillars of the *i*^*th*^ row and the (*i* − 1)^*th*^ row. *J* is the center-to-center distance between the pillars in any row. *T* is the center-to-center distance between the *i*^*th*^ row and the (*i* − 1)^*th*^ row.

The basic design of RAPID has been described in detail in an earlier manuscript [26]. We now provide detailed guidelines to the user for choosing various geometrical parameters of the device.

During the mask design, one should choose the value of the inlet diameter to be slightly larger than the size of the punch to give the user some room during punching. Otherwise, the first row of pillars might get damaged due to error in punching. For example, we used a biopsy punch of diameter 1.5 mm. Hence, in our design the inlet diameter was set to be 1.7 mm.

We used equation 1 to determine the number of pillars (*n*), the pillar diameter (*d*) and the gap (*g*) between pillars of zone 1.

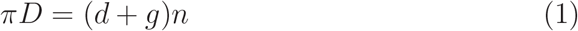

where,

*D* is twice the distance from center of the device to the center of the first row of pillars,

*d* is the diameter of the pillar,

*n* is the number of pillars in the first row,

*g* is the gap between the pillars.

Previously, we demonstrated the operation of RAPID with rigid particles. When using deformable objects (e.g., cells), both size and deformability need to be considered. Because of deformability the ‘effective size’ of a cell is always smaller than its actual size. Here, the value of ‘*g*’ should be smaller than the size of the cell to be blocked.

In our design, WBCs are captured by zone 1. The size range of WBCs is from 8 *μ*m to 30 *μ*m. Therefore, we chose *g* = 6*μ*m. If we choose a large pillar diameter (*d*), it is easy to fabricate during lithography. On the other hand, if ‘*d*’ is small, then there are more pillars (i.e. large *n*) in a row. This increases the number of the parallel paths for the cells and delays clogging. To balance these two considerations, we recommend the value of *d* to be 4*g*. Therefore, in our design *d* = 24 *μ*m. Once *d* and *g* are fixed, the number of pillars (*n*) is given by equation 2.

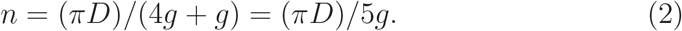

The pillar gap and the number of pillars should remain the same for all rows in a particular zone to keep the radial flow path constant. Therefore, the pillar diameter (*d*) has to increase as we move radially outwards, as given by equation 1. The increase in *D* (= *D*_2_ - *D*_1_) in moving from the first row to the second row is given by equation 3.

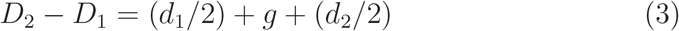

where *d*_1_ and *d*_2_ are the pillar diameters of the first and the second row respectively. Generalizing this formula for i rows we can write

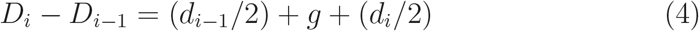

The ability to capture a cell increases with the increase in the number of rows. On the other hand, increasing the number of rows increases the device footprint, leading to a drop in the inlet pressure in the radial direction. Based on our previous experiments [26], we chose N = 9 because 99% of the beads were trapped by the first seven rows. Based on these considerations, the WBC capture zone had 180 pillars in each of the nine rows, with pillar diameters ranging from 24 *μ*m to 30 *μ*m, and a pillar gap of ~ 6 *μ*m.

The same design considerations are used to design zone 3. The RBC capture zone has 360 pillars in each of the five rows, with pillar size ranging from 24 *μ*m to 26 *μ*m, and 2 *μ*m pillar gap. We hypothesized that at low flow rates the 2 *μ*m pillar gap in the outermost cluster of pillars (RBC capture zone) would stop most of the RBCs from going through and would only allow the platelets to move radially outwards. The height of the device was reduced to 5 *μ*m (Supplementary figure S1) to prevent the biconcave RBCs from flipping on their sides and passing through gaps much smaller than their diameter.

The designing of zone 2 is slightly more complex because the sucessive rows of pillars are shifted by an angle *θ*. The pillar gap in zone 2 should be larger than the pillar gaps of zones 1 and 3 to let all cells through. The main purpose of this zone is to set up a cross flow in the device, facilitating long-term clog-free operation. The pillar diameter of the first row in zone 2 should be equal to the pillar diameter of the last row of zone 1. The radial distance between zones 1 and 2 is set to be equal to the pillar gap in last row of zone 1. Similarly, the radial distance between zones 2 and 3 is equal to the pillar gap in the last row of zone 2.

Next we use simple geometrical arguments to find a relationship (equation 5) between the angular displacement (*θ*), the pillar diameter (*d*), the pillar gap (*g*), the center-to-center distance (*T*_*i*−1_) between the *i*^th^ and the (*i* − 1)^th^ rows, and the center-to-center distance (*J*) of the pillars of particular row. The detailed calculation is given in the supplementary information. Equation 5 gives us the range of values for *θ*. As reported in our previous work [26], one can run FEM simulations for different values of theta and find out at what value the cross flow has the maximum strength.

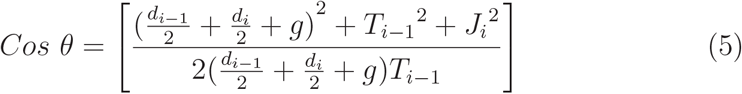

where,

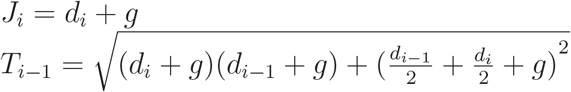

Accordingly, the consecutive rows of pillars in AD zone are shifted by 7*°*. The pillar gap varied from 8 *μ*m to 10 *μ*m over nine rows. There were 180 pillars in each row, with pillar sizes ranging from 32 *μ*m to 40 *μ*m. The angular displacement was expected to facilitate a cross flow in the AD zone.

**Figure 3:**
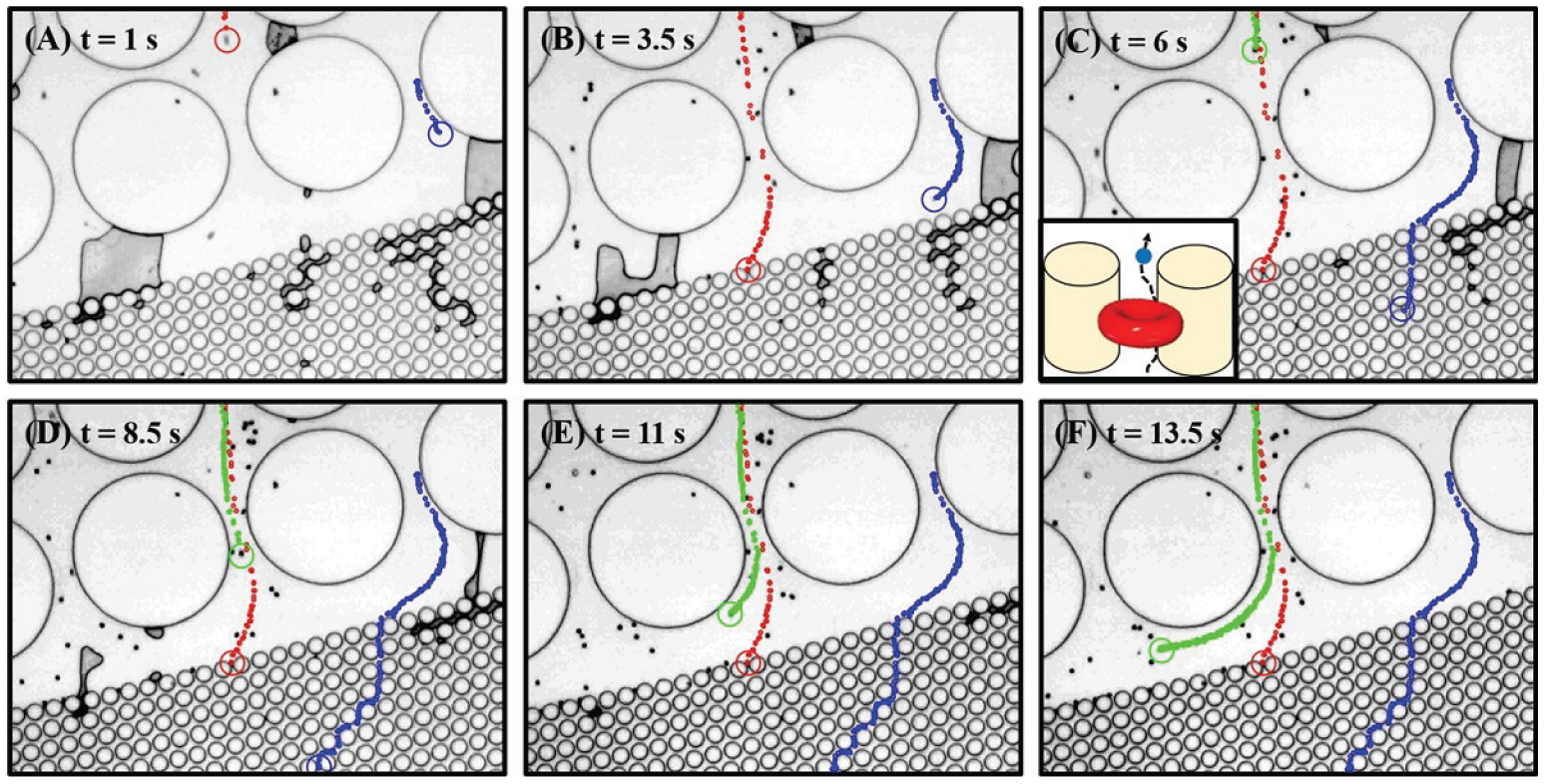
Time lapse images from the supplementary video SV2 showing the paths followed by RBCs and platelets in RAPID at different time points. (A-C) At the start of the experiment (t = 1 s to t = 6 s), RBCs and platelets follow a radial path (red track) to reach the RBC capture zone, where RBCs are stopped. The smaller cells can still follow a radial path (blue track) towards the platelet outlet. The schematic in the inset in panel C shows that there is room for platelets to pass through, even if an RBC is stuck flat between two pillars. (D) Around t = 8.5 s, a few of the radial paths are found to be blocked by the trapped RBCs. (E-F) This increases the strength of the cross flow and forces the other RBCs to follow the cross-flow (green track) towards the RBC outlet instead of taking the radial route.

### 3.2 Switching from dead-end to cross-flow operation

We first wished to check whether the RBCs move in a flat orientation in our device. The supplementary movie SV1 shows two RBCs approaching the RBC capture zone in RAPID chips with 10 *μ*m and 5 *μ*m heights respectively. In the 10 *μ*m height chip, the RBC flips on its side and passes through the 2 *μ*m pillar gaps. In contrast, the RBCs are oriented flat in the 5 *μ*m height chip. These flat RBCs can be stopped by the small pillar gaps at low flow rates.

We tested our hypothesis of flow switching using dilute blood so that the paths of RBCs and platelets can be clearly tracked. Fig. 3 contains some snapshots from the supplementary video SV2 and shows the paths taken by three different cells at different time points. Initially (panels A-D) RAPID functions like a dead-end pillar device, where RBCs (red track) and platelets (blue track) follow the radial path to reach the RBC capture zone. Here the RBCs are stopped flat at a low flow rate by the 2 *μ*m pillar gap and the 5 *μ*m height of the chamber. Since the platelets are much smaller than the RBCs, they can pass through (blue track) these gaps, towards the platelet outlet even when an RBC is sometimes stuck between two pillars (schematic shown in the inset in Fig. 3C). Once a few RBCs are stuck outside the RBC capture zone, the strength of the radial flow is reduced, and a cross flow towards the RBC outlet is strengthened (panels E-F). Newer RBCs (and some platelets) reaching this zone now follow the cross-flow (green track) towards the RBC outlet. It is the automatic switching from dead-end to a cross-flow operation that allows RAPID to function for several hours without additional buffer injection (like deterministic lateral displacement devices [27, 17]) or reverse flow (like dead-end filters [11]).

### 3.3 Operation of RAPID with whole blood

We next tested the operation of RAPID with whole blood in an experiment lasting 6h (supplementary video SV3). Blood was passed through the device at a flow rate of 2 *μ*l/min, and time-lapse images were acquired after every 4 sec. The initial images were taken using a 10X objective to visualize the flow pattern in the entire device, and the later images were taken using a 20X objective to focus on the region between the AD and the RBC capture zones. As hypothesized, the sample flow was radial in the initial stage (Fig. 4A). The strength of both radial and cross flows increased substantially at ~ 15 min, and the strong flows were maintained for ~ 4h. After this point, the flow was found to have slowed down. We noticed in the video that some RBCs were sticking to each other (forming rouleaux) and moving as larger objects through the obstacles. The sluggish flow continued until 6h when the experiment was stopped. The supplementary video SV3 shows that there were some small pockets in the device where the flow was sluggish throughout. Since there are multiple parallel radial paths in RAPID unlike dead-end devices, the presence of these zones did not affect the overall device operation.

**Figure 4:**
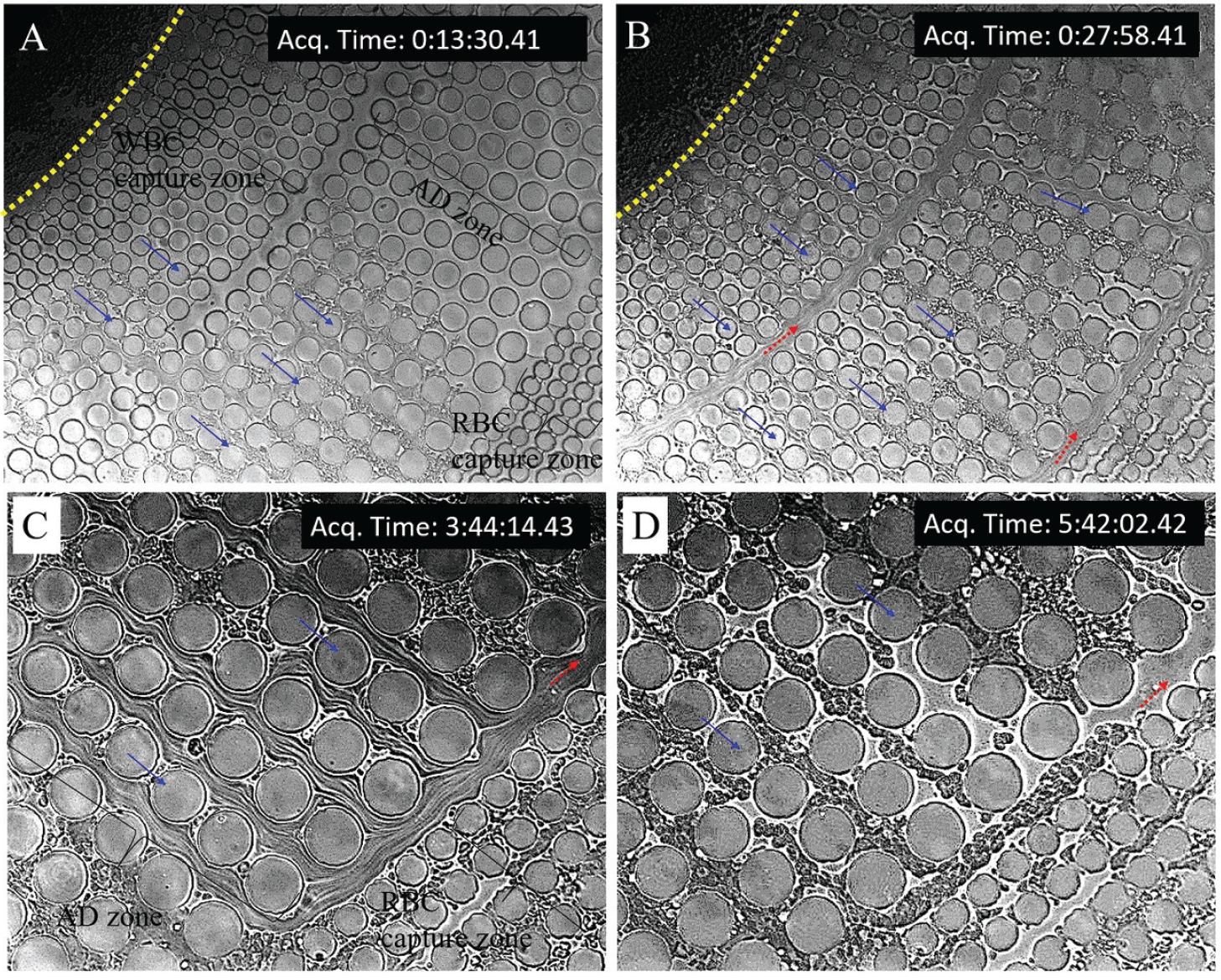
Time lapse microscope images acquired from the continuous operation of RAPID with whole blood for 6 h at a flow rate of 2 *μ*l/min. Panels A and B were captured using a 10X objective to record the sample flow over all three zones. The yellow dotted line indicates the inlet. Panels C and D were captured using a 20X objective to focus on the region between AD and RBC capture zones. (A) A snapshot taken at ~ 13 min shows that the sample flow is primarily in the radial direction (indicated by blue solid arrows). (B) An image acquired at ~ 28 min shows the presence of strong cross flows in the device. The cross-flow is the strongest in the regions between the successive zones, as indicated by the red dotted arrows. The angular displacement of the pillars in the AD zone keeps these flows unidirectional. (C) At 3h 44 min, strong radial and cross flows are still maintained in the device. The streaks seen in the image, instead of the individual cells, indicate the high speed of the cells contained in these flows. (D) At 5h 42 min, both the radial and the cross-flow are found to have slowed down. At this point, primarily the plasma (and the platelets therein) continue to flow. The experiment was stopped after 6h.

**Figure 5:**
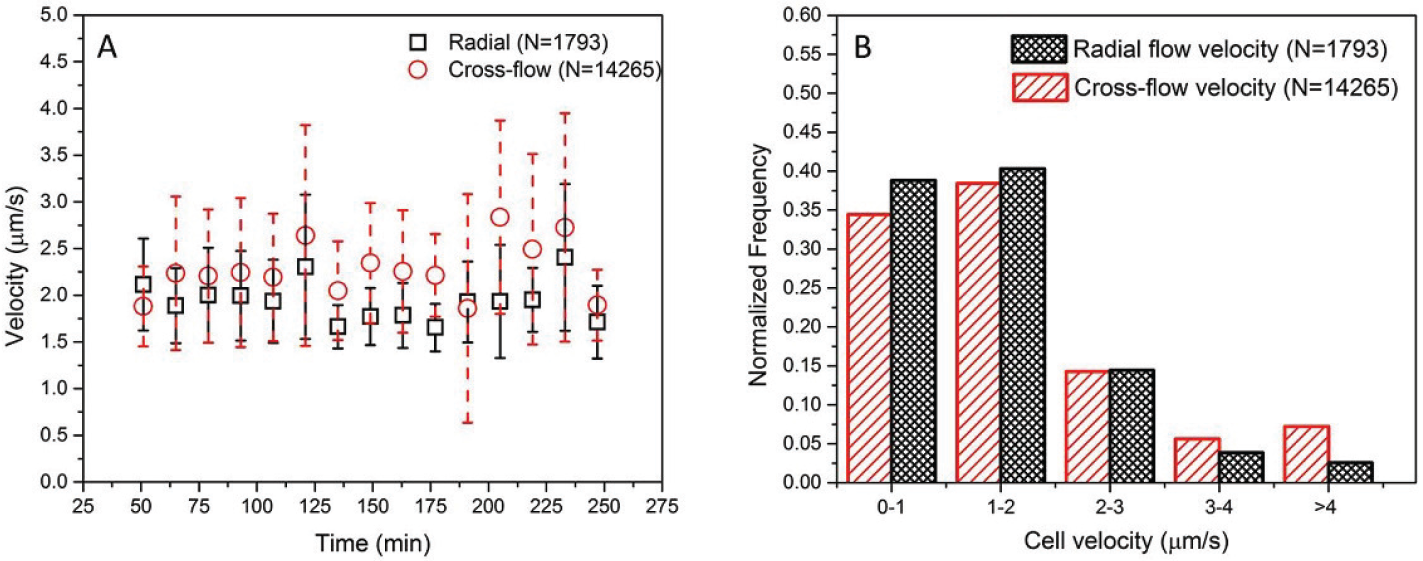
Velocity profile of the cells in radial and cross flows over the duration of the experiment shown in video SV4. The velocity of the cells was tracked from 1 hour after starting the experiment until ~ 4 hours. (A) Most of the time, the cells in cross flow had a higher mean velocity compared to the cells in radial flow. The cells continued to be in motion until the end of the experiment. The error bars indicate the standard deviation. *N* indicates the total number of cells analyzed. (B) The normalized histogram shows the velocity distribution of the cells in radial and cross flow for the entire duration of the experiment. About 72% cells in the cross flow and 79% cells in the radial flow had a velocity less than 2 *μ*m/s.

As seen in Fig. 4 and SV3, some of the RBCs manage to squeeze through the RBC capture zone and reach the platelet outlet at a flow rate of 2 *μ*l/min. We also repeated this experiment at a lower flow rate of 600 nl/min for close to 4h (supplementary video SV4). The number of RBCs entering the outermost zone is much lower when the flow rate is low (600 nl/min). Hence, an increase in the flow rate lowers the purity of the platelets collected from RAPID. This observation is discussed in more detail in the next section.

Figure 5 shows the velocity profile of the cells in radial and cross flows in order to quantify our observations in the video SV4. For the first one hour, the video was acquired with lower magnification and it was difficult to extract the individual cell tracks. Therefore, the velocity was tracked from 1 hour after starting the experiment until ~ 4 hours. Non-zero radial and cross flow velocity throughout the experiment justified our claim of clog-free operation of RAPID with whole blood. Most of the time, the cells in cross flow had a slightly higher velocity compared to the cells in radial flow. As expected, the number of cells moving in the cross flow was almost 10 times higher than the number of cells moving in the radial flow. As seen in the videos (SV3 and SV4), sometimes a few cells clumped together and slowed down the cross flow. The cross flow velocity suddenly increased when these clumps got cleared from time to time. This led to a larger variation in the cross flow velocity throughout the experiment.

We found that there are two important requirements when handling whole blood in RAPID for a long duration. First, the RBCs in the inlet tube tend to settle over time. So, a vibrator motor was attached to the inlet tube to periodically resuspend the cells. Second, the connectors and the tubing sometimes came out of the PDMS chip under the strong vibration of the motor and weight of the connectors. Therefore, these needed to be reinforced, as described in the supplementary information. These measures were not necessary when we worked with dilute blood samples.

### 3.4 Platelet enrichment from whole blood

We performed platelet enrichment experiments in RAPID as an illustration of the ability of the device to work with whole blood. We calculated separation purity, recovery, enrichment factor and the throughput of platelet outlet to characterize these experiments. We did not consider the contribution of WBCs while calculating these parameters. This is because the majority of the WBCs were prevented from entering the device at the inlet itself by the 6*μ*m pillar gaps and the 5*μ*m device height. The large inlet area (1.5 mm diameter) allowed us to avoid clogging of the device due to the accumulation of WBCs. The experiments were continued until approximately 50 *μ*l volume of sample was collected from each of RBC and platelet outlets for analysis using a hematology analyzer.

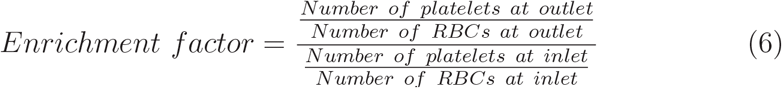

Equation 6 defines the enrichment factor of platelets in RAPID. It is a measure of how well the desired cells (platelets) are concentrated compared to the undesired cells (RBCs in this case) at the platelet outlet. Fig. 6A confirms that RAPID can concentrate platelets from whole blood up to 60 times at the lowest flow rate of 600 nl/min. Throughput (Fig. 6B) measures the average volume of sample collected from the platelet outlet in a minute. The highest throughput of platelet collection is ~ 800 nl/min. Recovery (Fig. 6C) is a measure of the number of platelets lost in the device during the experiment. Recovery is given by the ratio of the number of platelets at all outlets to the number of platelets at the inlet and remains ~ 95% at all flow rates. The separation purity indicates what percentage of the cells (RBCs and platelets) collected from the platelet outlet are actually platelets. As seen from Fig. 6D, the separation purity of platelets decreases with an increase in flow rate. We achieved a maximum of ~ 70% purity at the lowest flow rate of 600 nl/min. All data are plotted as histograms with the mean value, and the standard deviation indicated.

**Figure 6:**
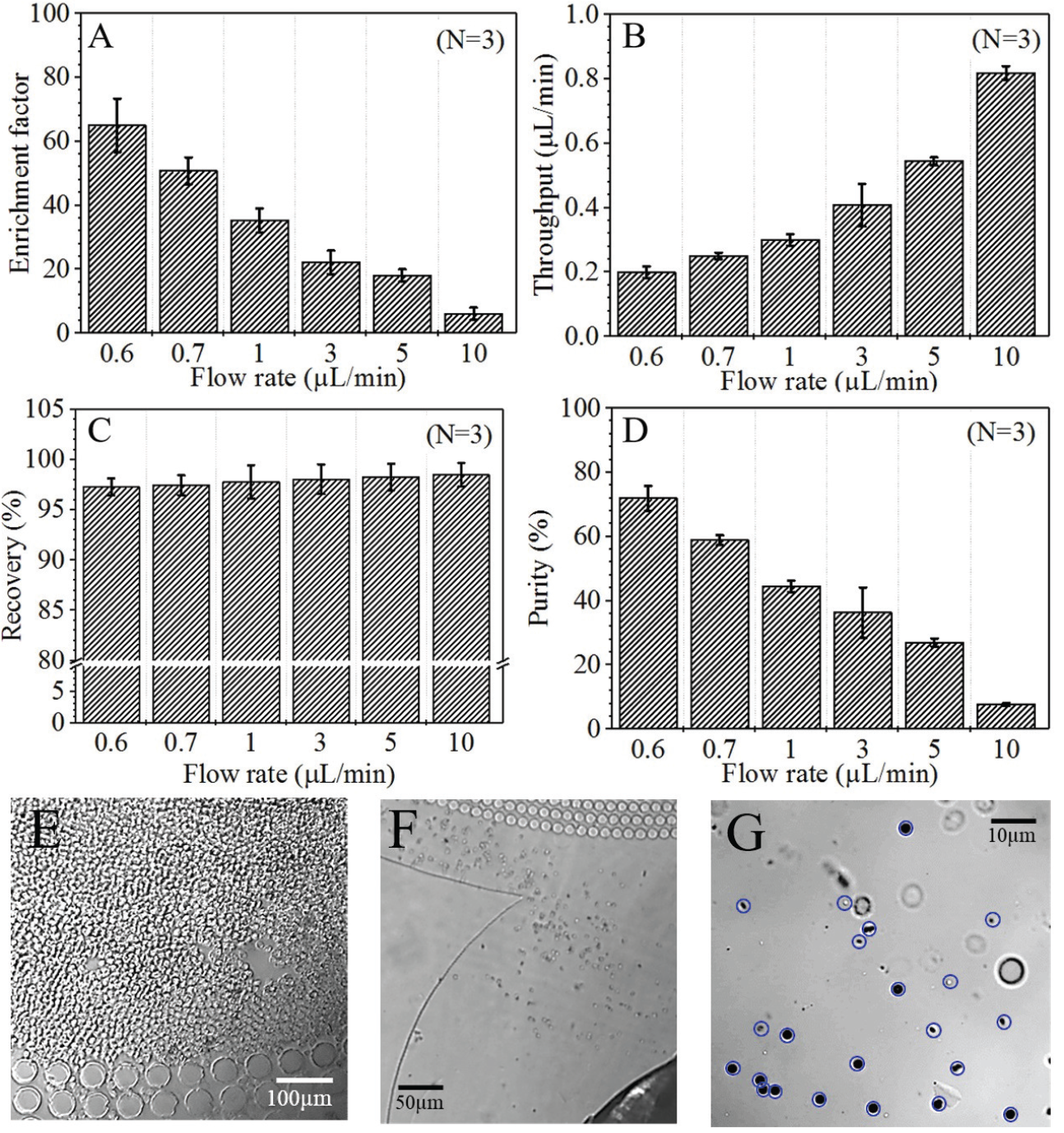
Platelet separation parameters as a function of flow rate in RAPID. (A) The platelet enrichment factor decreases with increase in flow rate. We can achieve 60-fold enrichment at the lowest flow rate of 600 nl/min. (B) In contrast, throughput at platelet outlet increases with an increase in the flow rate. (C) The recovery remains unaffected by the flow rate variation. (D) The separation purity of platelets decreases from 70% to 10% with an increase in flow rate from 600 nl/min to 10 *μ*l/min. As the pressure is increased, more and more deformable RBCs squeeze through the RBC capture zone to reach the platelet outlet. (E) A microscope image of the inlet (taken using 20X objective) shows a large number of RBCs. (F) A microscope image of the platelet outlet (taken using 10X objective) shows that there are very few RBCs present in comparison with the inlet. Images (E) and (F) were taken at a flow rate of 600 nl/min. (G) An image of the platelets (blue circles) present at the outlet. This image was acquired using a 63X (1.4 NA) objective.

There is a trade-off between the different performance parameters as a function of the inlet flow rate. Both platelet purity and enrichment peak at the lowest flow rate. On the other hand, throughput improves at higher flow rates. At lower flow rates, most RBCs are stopped by the pillar gaps of the RBC capture region and follow the cross flow to the RBC outlet. As the flow rate is increased, more and more deformable RBCs are squeezed through the gaps towards the platelet outlets. The supplementary videos SV3 and SV4 confirm this observation. We need to strike a balance between different performance measures when choosing an optimal flow rate.

Panels 6E-G show some images of the cells taken at the inlet and the platelet outlet after running an experiment at 600 nl/min inlet flow rate. The inlet image (panel E) shows that there are too many RBCs entering the device. The image of the platelet outlet (panel F) confirms that very few RBCs make it to the platelet outlet at low flow rates. Panel G shows an image of the platelet outlet taken using a 63X objective (1.4 NA). The blue circles indicate the platelets. Since, there are approximately 20 RBCs for every platelet in whole blood, an enrichment factor more than 20 can help to offset the effect of RBCs during platelet count. Such images, along with a high enough enrichment factor, can potentially be used to obtain the platelet count from whole blood samples without lysing cells.

### 3.5 Comparison with similar passive microfluidic devices

We finally compared (table 1) how RAPID handles whole blood in comparison to passive microfluidic devices for plasma or platelet separation. We looked at the performance metrics, such as, the operation time of the device, the volume of blood handled during operation, sample recovery, and the throughput at the desired sample outlet. None of the obstacle-based passive devices (e.g. membrane or plug filtration, trench, weir, etc.) reported platelet separation from whole blood. Therefore, we also included two hydrodynamic devices (e.g. elastoinertial and hydrophoretic) which reported platelet separation from dilute blood, in our comparison.

**Table 1:** Comparison of passive microfluidic devices focused on platelet and plasma separation from whole blood

| Technique | Working time (min) | Volume handled (μl) | Recovery (%) | Throughput (μl/min) | Output Sample |
| --- | --- | --- | --- | --- | --- |
| Membrane filtration [7] | 1 | 50 | 30 | 6.5 | Plasma |
| Membrane filtration [6] | 20 | 5 | 20 | - | Plasma |
| Plug filtration [8] | 10 | 10 | 1.9 | 0.02 | Plasma |
| Trench [21] | 10 | 5 | - | 0.8 | Plasma |
| Weir [20] | 3 | - | - | 0.8 | Plasma |
| Elasto-inertial [28] | - | - | 92 | 0.5 | Platelets |
| Hydrophoretic [29] | - | - | 41 | 2 | Platelets |
| **RAPID** | **360** | **720** | **95** | **0.8** | **Platelets** |

All the obstacle-based devices reported operation times of 20 min or less, and sample volumes of 50 *μ*l or less. In contrast, RAPID can operate up to ~ 6h by continuously removing the RBCs through the cross flow, and can handle more than 700 *μ*l of whole blood. The sample recovery in RAPID is ~ 95%. It is comparable to that achieved by the elastoinertial effect, and much larger than the recovery obtained by filtration. Chung *et al.* have reported a throughput of > 6 *μ*l/min using membrane filtration. Most filtration devices achieve throughput of <1 *μ*l/min, comparable to the throughput achieved in RAPID for platelet separation. It should be noted that the entire device throughput (i.e. the combined throughput of RBC and platelet outlets) is much higher.

## 4 Conclusion

Handling large volumes of whole blood for a long duration in passive microfluidic pillar-based devices is extremely challenging. Most reported devices use diluted blood samples to prevent clogging of the device. Here we adapted the design of a previously-reported <u>ra</u>dial <u>pi</u>llar device (RAPID) for continuous handling of concentrated suspensions of deformable objects, such as, whole blood. We demonstrated the operation of RAPID with whole blood for up to 6 h at a flow rate of 2 *μ*l/min. We have also provided detailed guidelines to design RAPID for any application. As an illustration of the ability to work with whole blood, we performed platelet enrichment in our device, and achieved an enrichment factor of 60X at an inlet flow rate of 600 nl/min. Due to its capability to handle large volumes of whole blood samples for several hours, RAPID can potentially be used for many different applications, such as, rare cell capture, studying the effect of drugs in real-time, cell migration, etc.

## 5 Acknowledgements

We acknowledge funding for cleanroom access from the Centre for Nano-electronics Phase 2 project (MeiTY, Govt. of India). All lithography was performed in the clean-room of Indian Institute of Technology Bombay’s nanofabrication facility. We thank the LSM confocal microscope facility housed in the Department of Biosciences and Bioengineering for aquiring the images in figure 6. We acknowledge financial (travel) support from the Wadhwani Research Centre for Bioengineering. Finally, we thank Kaushalya Foundation Medical Trust Hospital for technical support and discussions.

## References

[1] K. Kamei, K. Tajima, N.T. Huy, T. Kariu, et al., One-step concentration of malarial parasite-infected red blood cells and removal of contaminating white blood cells, Malaria Journal 3(1), 7 (2004)

[2] R. Mariella, Sample preparation: the weak link in microfluidics-based biodetection, Biomedical microdevices 10(6), 777 (2008)

[3] H.M. Ji, V. Samper, Y. Chen, C.K. Heng, T.M. Lim, L. Yobas, Silicon-based microfilters for whole blood cell separation, Biomedical microdevices 10(2), 251 (2008)

[4] S. Thorslund, O. Klett, F. Nikolajeff, K. Markides, J. Bergquist, A hybrid poly (dimethylsiloxane) microsystem for on-chip whole blood filtration optimized for steroid screening, Biomedical microdevices 8(1), 73 (2006)

[5] J. Moorthy, D.J. Beebe, In situ fabricated porous filters for microsystems, Lab on a Chip 3(2), 62 (2003)

[6] D.S. Lee, Y.H. Choi, Y.D. Han, H.C. Yoon, S. Shoji, M.Y. Jung, Construction of membrane sieves using stoichiometric and stress-reduced si 3 n 4/sio 2/si 3 n 4 multilayer films and their applications in blood plasma separation, ETRI Journal 34(2), 226 (2012)

[7] K.H. Chung, Y.H. Choi, J.H. Yang, C.W. Park, W.J. Kim, C.S. Ah, G.Y. Sung, Magnetically-actuated blood filter unit attachable to premade biochips, Lab on a Chip 12(18), 3272 (2012)

[8] C. Li, C. Liu, Z. Xu, J. Li, Extraction of plasma from whole blood using a deposited microbead plug (dmbp) in a capillary-driven microfluidic device, Biomedical microdevices 14(3), 565 (2012)

[9] Y. Cheng, X. Ye, Z. Ma, S. Xie, W. Wang, High-throughput and clogging-free microfluidic filtration platform for on-chip cell separation from undiluted whole blood, Biomicrofluidics 10(1), 014118 (2016)

[10] V. VanDelinder, A. Groisman, Separation of plasma from whole human blood in a continuous cross-flow in a molded microfluidic device, Analytical chemistry 78(11), 3765 (2006)

[11] S.M. McFaul, B.K. Lin, H. Ma, Cell separation based on size and deformability using microfluidic funnel ratchets, Lab on a chip 12(13), 2369 (2012)

[12] Y. Yoon, S. Kim, J. Lee, J. Choi, R.K. Kim, S.J. Lee, O. Sul, S.B. Lee, Clogging-free microfluidics for continuous size-based separation of microparticles, Scientific reports 6, 26531 (2016)

[13] X. Chen, C.C. Liu, H. Li, et al., Microfluidic chip for blood cell separation and collection based on crossflow filtration, Sensors and Actuators B: Chemical 130(1), 216 (2008)

[14] Z. Geng, Y. Ju, W. Wang, Z. Li, Continuous blood separation utilizing spiral filtration microchannel with gradually varied width and micropillar array, Sensors and Actuators B: Chemical 180, 122 (2013)

[15] E. Sollier, H. Rostaing, P. Pouteau, Y. Fouillet, J.L. Achard, Passive microfluidic devices for plasma extraction from whole human blood, Sensors and Actuators B: Chemical 141(2), 617 (2009)

[16] K. Aran, A. Fok, L.A. Sasso, N. Kamdar, Y. Guan, Q. Sun, A. Ündar, J.D. Zahn, Microfiltration platform for continuous blood plasma protein extraction from whole blood during cardiac surgery, Lab on a Chip 11(17), 2858 (2011)

[17] D.W. Inglis, N. Herman, G. Vesey, Highly accurate deterministic lateral displacement device and its application to purification of fungal spores, Biomicrofluidics 4(2), 024109 (2010)

[18] J. McGrath, M. Jimenez, H. Bridle, Deterministic lateral displacement for particle separation: a review, Lab on a Chip 14(21), 4139 (2014)

[19] K.K. Zeming, T. Salafi, C.H. Chen, Y. Zhang, Asymmetrical deterministic lateral displacement gaps for dual functions of enhanced separation and throughput of red blood cells, Scientific reports 6, 22934 (2016)

[20] T. Tachi, N. Kaji, M. Tokeshi, Y. Baba, Simultaneous separation, metering, and dilution of plasma from human whole blood in a microfluidic system, Analytical chemistry 81(8), 3194 (2009)

[21] I.K. Dimov, L. Basabe-Desmonts, J.L. Garcia-Cordero, B.M. Ross, A.J. Ricco, L.P. Lee, Stand-alone self-powered integrated microfluidic blood analysis system (simbas), Lab on a Chip 11(5), 845 (2011)

[22] M. Kersaudy-Kerhoas, E. Sollier, Micro-scale blood plasma separation: from acoustophoresis to egg-beaters, Lab on a Chip 13(17), 3323 (2013)

[23] V. Vickerman, J. Blundo, S. Chung, R. Kamm, Design, fabrication and implementation of a novel multi-parameter control microfluidic platform for three-dimensional cell culture and real-time imaging, Lab on a Chip 8(9), 1468 (2008)

[24] D. Wlodkowic, J.M. Cooper, Tumors on chips: oncology meets microfluidics, Current opinion in chemical biology 14(5), 556 (2010)

[25] S. Nagrath, L.V. Sequist, S. Maheswaran, D.W. Bell, D. Irimia, L. Ulkus, M.R. Smith, E.L. Kwak, S. Digumarthy, A. Muzikansky, et al., Isolation of rare circulating tumour cells in cancer patients by microchip technology, Nature 450(7173), 1235 (2007)

[26] N. Mehendale, O. Sharma, C. D’Costa, D. Paul, A radial pillar device (rapid) for continuous and high-throughput separation of multi-sized particles, Biomedical Microdevices 20(1), 6 (2018)

[27] J.A. Davis, D.W. Inglis, K.J. Morton, D.A. Lawrence, L.R. Huang, S.Y. Chou, J.C. Sturm, R.H. Austin, Deterministic hydrodynamics: taking blood apart, Proceedings of the National Academy of Sciences 103(40), 14779 (2006)

[28] J. Nam, H. Lim, D. Kim, H. Jung, S. Shin, Continuous separation of microparticles in a microfluidic channel via the elasto-inertial effect of non-newtonian fluid, Lab on a Chip 12(7), 1347 (2012)

[29] S. Choi, T. Ku, S. Song, C. Choi, J.K. Park, Hydrophoretic high-throughput selection of platelets in physiological shear-stress range, Lab on a Chip 11(3), 413 (2011)

